# Effects of Repetitive Mild Traumatic Brain Injury on Corticotropin-Releasing Factor Modulation of Lateral Habenula Excitability and Motivated Behavior

**DOI:** 10.1101/2024.04.16.589760

**Authors:** William J. Flerlage, Sarah C. Simmons, Emily H. Thomas, Shawn Gouty, Mumeko C. Tsuda, T. John Wu, Regina C. Armstrong, Brian M. Cox, Fereshteh S. Nugent

## Abstract

Mild traumatic brain injury (mTBI) is a significant health burden due to mTBI-related chronic debilitating cognitive and psychiatric morbidities. Recent evidence from our laboratory suggests a possible dysregulation within reward/motivational circuit function at the level of a subcortical structure, the lateral habenula (LHb), where we demonstrated a causal role for hyperactive LHb in mTBI-induced motivational deficits in self-care grooming behavior in young adult male mice when exposed to mTBI injury during late adolescence (at ∼8 weeks old). Here we extended this observation by further characterizing neurobehavioral effects of this repetitive closed head injury model of mTBI in both young adult male and female mice on LHb excitability, corticotropin releasing factor (CRF) modulation of LHb activity, and behavioral responses of motivation to self-care behavior, and approach versus avoidance behavior in the presence of a social- or threat-related stimulus. We show that mTBI increases LHb spontaneous tonic activity in female mice similar to what we previously observed in male mice as well as promoting LHb neuronal hyperexcitability and hyperpolarization-induced LHb bursting in both male and female mice. Interestingly, mTBI only increases LHb intrinsic excitability in male mice coincident with higher levels of the hyperpolarization-activated cation currents (HCN/Ih) and reduces levels of the M-type potassium currents while potentiating M-currents without altering intrinsic excitability in LHb neurons of female mice. Since persistent dysregulation of brain CRF systems is suggested to contribute to chronic psychiatric morbidities and that LHb neurons are highly responsive to CRF, we then tested whether LHb CRF subsystem becomes engaged following mTBI. We found that *in vitro* inhibition of CRF receptor type 1 (CRFR1) within the LHb normalizes mTBI-induced enhancement of LHb tonic activity and hyperexcitability in both sexes, suggesting that an augmented intra-LHb CRF-CRFR1-mediated signaling contributes to the overall LHb hyperactivity following mTBI. Behaviorally, mTBI diminishes motivation for self-care grooming in female mice as in male mice. mTBI also alters defensive behaviors in the looming shadow task by shifting the innate defensive behaviors towards more passive action-locking rather than escape behaviors in response to an aerial threat in both male and female mice as well as prolonging the latency to escape responses in female mice. While, this model of mTBI reduces social preference in male mice, it induces higher social novelty seeking during the novel social encounters in both male and female mice. Overall, our study provides further translational validity for the use of this preclinical model of mTBI for investigation of mTBI-related reward circuit dysfunction and mood/motivation-related behavioral deficits in both sexes while uncovering a few sexually dimorphic neurobehavioral effects of this model that may differentially affect young males and females when exposed to this type of mTBI injury during late adolescence.

## Introduction

Traumatic brain injury (TBI) remains a leading cause of death and disability in the United States with profound economic and societal impact. Specifically, mild TBI (mTBI) characterized by transient alteration of consciousness comprises the majority of TBI. mTBI is a significant health burden due to an increased likelihood of long-lasting impairments in cognition, mood/emotional regulation, social interactions and risk-taking behaviors in susceptible individuals^1, 2^. mTBI-related negative mood outcomes include affective deficits (e.g., depression, apathy), emotional dysregulation and social dysfunction (irritability, aggression, suicidality, social withdrawal, anxiety) and posttraumatic stress disorder (PTSD, heightened fear and/or freezing responses to threats, avoidance, hypervigilance)^3–6^. Concussive injury as a subtype of mTBI occurs during the transfer of a mechanical energy to the brain following an external force to the head in the absence of any structural brain damages^7^. The risk of chronic mood-related psychiatric morbidity following mTBI also increases with repeated mTBI as seen in high-risk populations such as contact-sports athletes, military personnel, and victims of intimate partner violence ^2, 8, 9^. Moreover, accumulating evidence suggests sex differences in long-term negative outcomes of mTBI; specifically females at certain ages (from ∼ 35 to 50 years of age) are at higher risk of worse mood outcomes including depression and anxiety due to declines in the neuroprotective contributions of estrogen and progesterone, and age-related increased vulnerability to various environmental stressors (though there are also sex differences in reporting biases that should be accounted) ^10–15^. Recently, emerging evidence has suggested the lateral habenula (LHb), an anti-reward brain region implicated in motivation and decision making, as a common and major neural substrate underlying higher susceptibility for development of negative affective symptoms in depression and mood disorders^16–19^. LHb promotes avoidance behaviors in response to aversive and unpleasant events or the unexpected omission of reward through integration of the information from forebrain limbic structures and conveying an “anti-reward” signal through suppression of ventral tegmental area (VTA) dopamine and dorsal raphe nucleus serotonin systems ^17, 20, 21^. LHb neurons provide glutamatergic projections to the substantia nigra, VTA, rostromedial tegmental area (RMTg), dorsal raphe nucleus, locus coeruleus and periaqueductal gray and receive glutamatergic, GABAergic and co-releasing glutamate/GABA inputs from the basal ganglia and diverse limbic areas such as medial prefrontal cortex (mPFC), entopeduncular nucleus, and lateral hypothalamus ^17, 20, 21^. In general, LHb hyperactivity is a common finding in negative affective states including anhedonia, lack of motivation and social withdrawal, the hallmarks of reward deficits and depression ^22–25^. Not surprisingly, in the last decade LHb has gained significant attention as a critical anatomical therapeutic target for identification and development of new, fast-acting, and more effective antidepressants ^26–28^ and circuit-based neuromodulation ^29–32^. Yet an understanding of reward- and motivation-related LHb dysfunction in mTBI-induced psychiatric morbidities in both biological sexes remains elusive. This is a significant knowledge gap given that mTBI-related psychopathologies are often treatment-resistant ^33–35^. For example, while classical antidepressants such as serotonin reuptake inhibitors (SSRIs) were initially reported to be effective in relieving mTBI-related depression ^36, 37^, recent evidence highlighted that SSRIs were no more effective than placebo in people with depression following a TBI ^33–35^. This may suggest differences in pathophysiology and antidepressant responses of TBI-induced depression from non-TBI forms of depression. Given the significant lack of preclinical mTBI studies in females as well as studies focused on mTBI-related reward circuit dysfunction in both biological sexes^15, 38^, we have developed a preclinical model of mTBI using repetitive closed injury mouse model of mTBI which induces behavioral deficits in motivational self-care grooming in sucrose splash test ^39^ and social interaction test ^40^ as well as persistent LHb hyperactivity through a shift in synaptic excitation and inhibition (E/I) balance toward excitation, and via plasma membrane insertion of calcium-permeable (CP) AMPARs in LHb neurons in male mice ^39^. We demonstrated that limiting LHb hyperactivity by chemogenetic inhibition of LHb neurons was sufficient to reverse mTBI-induced delays in self-care grooming behavior in male mice supporting a causal link between LHb hyperactivity and mTBI-induced self-grooming deficits^39^.

TBI patients also experience neuroendocrine dysfunction ^41–44^ with dysregulation of the hypothalamic pituitary axis (HPA) stress neuromodulator, corticotropin releasing factor (CRF), that has significant impact on stress neuronal responses and affective states following mTBI ^45–49^. Our previous studies have demonstrated that the rodent LHb is highly responsive to CRF^50, 51^. Earlier we reported that CRF acts through CRF receptor 1 (CRFR1)-protein kinase A (PKA) signaling, resulting in LHb hyperexcitability through PKA-dependent suppression of small conductance potassium SK channel activity, as well as presynaptic GABA release via retrograde endocannabinoid (eCB)-CB1 receptor signaling without altering glutamatergic activity in rat LHb neurons ^50^. Recently, we examined neuromodulatory effects of CRF on the mouse LHb excitability where we found that CRF exerted similar effects on mouse LHb with increased LHb intrinsic excitability coincident with higher input resistance, reduced levels of mAHPs and more negative AP thresholds. However, we observed that CRF uniformly attenuates GABAergic and glutamatergic transmission across all mouse LHb neurons^51^. Altogether, these findings suggest that CRF generally promotes LHb intrinsic excitability in rodent LHb, however it was unclear whether mTBI alters endogenous CRF/CRFR1 signaling tone which could then contribute to LHb hyperexcitability in mTBI male and female mice.

Here we further characterized neurobehavioral changes associated with this model where we observed that this model of mTBI results in similar alterations in LHb excitability, CRF-regulation of LHb activity and behavioral responses to sucrose splash and looming threat-related stimuli in male and female mice (with a few sex differences in intrinsic plasticity and social deficits), further supporting the translational utility of this mTBI model in both sexes for investigation of reward/motivational circuit dysfunction relevant to mTBI-related depression, anxiety and PTSD in humans.

## Materials and Methods

### Animals

All experiments were carried out in accordance with the National Institutes of Health (NIH) *Guide for the Care and Use of Laboratory Animals* and were approved by the Uniformed Services University Institutional Animal Care and Use Committee. C57BL/6 male mice (JAX) were acquired at ∼postnatal day 35-49 (PN35-P49) and allowed at least 72hr of acclimation before the initiation of any experimental procedures. Mice were group housed in standard cages under a 12hr/12hr light-dark cycle with standard laboratory lighting conditions (lights on, 0600-1800, ∼200lux) with ad libitum access to food and water except for social interaction and looming shadow task. A week before behavioral testing for social interaction test followed by the looming shadow task, mice were maintained under reversed light-dark cycle (lights off, 0600-1800) and individually housed 2 days prior to social interaction tests. All procedures were conducted beginning 2–4hr after the start of the light-cycle, unless otherwise noted. All efforts were made to minimize animal suffering and reduce the number of animals used throughout this study.

### Repetitive mild traumatic brain injury model

Beginning at ∼ PN56, mice were subjected to either repeated sham or repeated closed head injury (CHI) delivered by the Impact One, Controlled Cortical Impact (CCI) Device (Leica; Wetzler, Germany) utilizing parameters which were previously described^39, 40^. Mice were anesthetized with isoflurane (3.5% induction/2% maintenance) and fixed into a stereotaxic frame. Specifically, repeated CHI-CCI (mTBI group) consists of 5 discrete concussive impacts to the head delivered at 24hr intervals generated by an electromagnetically driven piston (4.0m/s velocity, 3mm impact tip diameter, a beveled flat tip, 1.0 mm depth; 200 ms dwell time) targeted to bregma as visualized through the skin overlying the skull following depilation. Sham surgery consisted of identical procedures without delivery of impact. Body temperature was maintained at 37^⸰^C throughout by a warming pad and isoflurane exposure and surgery duration was limited to no more than 5 minutes. Following sham or CHI-CCI surgery completion, mice were immediately placed in a supine position in a clean cage on a warming pad and the latency to self-right was recorded.

### Sucrose splash test

We performed behavioral testing in separate cohorts of mice for sucrose splash test followed by social interaction and then visual looming shadow tests or sucrose splash test followed by social interaction and then elevated zero maze separated by ∼2-7 days. Sucrose splash test was performed at 10-12 days following the final mTBI or sham procedure. Mice were video monitored throughout the sucrose splash test. Mice were individually introduced to an empty (7×11.5×4.5 inches) clear polycarbonate cage. Following a 10-min baseline assessment of behavioral activity the animal was gently removed from the testing arena, sprayed twice with an atomizer containing 10% sucrose solution onto the dorsal coat, returned to the test arena, and monitored for an additional 5 min. The 10% sucrose solution is a sticky substance that soils the animal’s coat, with the typical response being rapid initiation of vigorous grooming behaviors. Video recordings were assessed by an experimenter blinded to the condition of the subjects and scored for total grooming behavior and the latency to initiate the first bout of grooming after sucrose splash. Grooming is considered any movements involving active touching, wiping, scrubbing, or licking of the face, forelimbs, flank, or tail for greater than 3 consecutive seconds.

### Social interaction test

Three-chamber social interaction test with three-chamber sociability test apparatus (Stoelting Co., Wood Dale, IL) was used to evaluate social deficits. In the first session, the subject mouse was habituated to the three-chamber box for 10 min and then was placed in the middle chamber and the interaction times of the test mouse with either a novel age/sex-matched conspecific (the first stranger mouse placed at one of the chambers at one end of the arena) versus the other side chamber that was empty cup were recorded over 10 min. The movement of the test mouse was video recorded to determine the time related to social approach (defined as movement toward, circling, or sniffing of the stranger mouse in the social chamber). To test for social novelty or preference, we then placed a second stranger mouse inside an identical containment cup in the opposite side chamber that was the empty cup during the first session. Therefore, a first stranger mouse represents a familiar mouse that the test mouse has the chance to interact with for 10 min during the first session. We video recorded and monitored the same parameters described above in the second 10 min session to differentiate the behaviors between the test mouse in the presence of a first conspecific stranger (familiar) compared with a second conspecific stranger mouse (novel). In this and the following behavioral assays, the apparatus was always wiped clean with 70% ethanol between subjects and sanitized with Sanicloth wipes upon completion of tests.

### Elevated zero maze

The test evaluates anxiety and risk-taking behavior; mice are free to explore an elevated ring-shaped arena with two open and two closed quadrants (with elevated walls) and the number of entries and time spent in open and closed arms will be recorded over 10 min. The maze consisted of four arms (5 cm × 30 cm), including two closed arms having 20-cm high walls and two arms left open (open arms). The maze was elevated 40 cm above the floor. A camera above the maze relayed animal position information to Any-Maze software (Stoelting Co.), which reported distance traveled and relative amounts of time spent in the open and closed arms of maze.

### Looming-shadow test

The test mimics a predator threat for mice; mice adopt action-locking freezing (passive) or escape (active) defensive behaviors. The testing area (17 × 10 × 8.5 inches plastic arena) contains a 21-inch LCD monitor positioned above the arena facing downward that presents a visual stimulus. We placed a shelter (6×6×3 inches) in one side of area where the mouse can escape and hide underneath of it. The camera was placed in the lateral top part of the arena to record the behavior. Mice were habituated to the arena for 2 days for 10min/day. The mouse was placed in the arena for 5min habituation period on test day before presentation of visual stimulus. The mouse was presented by five overhead looming visual stimuli (a 2-cm black disk that was expanded to 20 cm in three distinct phases for a total of 9s) with at least 1 min inter-trial interval. The disk was present for 2 s, then expanded to 20 cm in 5s and then remained stable at this full size for an additional 2s. We discriminated the passive and active defensive behaviors based on the escape (active) to the shelter or the absence of any movement or repeated, discontinuous bouts of freezing during the trial as a passive action-locking/freezing behavior. The exclusion criterion that classified a mouse as a non-responder was when the mouse did not show any discernable response to the looming stimulus (no freezing or show any obvious change in ongoing behavior or while tried to escape, failed to get to the shelter within the 8s trial). We video recorded and blindly scored the type of defensive behavior (action-locking/escape) and latency to the escape behavior.

### Slice preparation

All electrophysiological experiments were performed at ∼4 weeks post-mTBI. Mice were deeply anesthetized with isoflurane and immediately transcardially perfused with ice-cold artificial cerebrospinal fluid (ACSF) containing (in mM): 126 NaCl, 21.4 NaHCO_3_, 2.5 KCl, 1.2 NaH_2_PO_4_, 2.4 CaCl_2_, 1.00 MgSO_4_, 11.1 glucose, 0.4 ascorbic acid; saturated with 95% O_2_-5% CO_2_. Brain tissue was kept on ice-cold ACSF and tissue sections containing LHb were sectioned at 220μm using a vibratome (Leica; Wetzler, Germany) and subsequently incubated in ACSF at 34 °C for at least 1-hr prior to electrophysiological experiments. For patch clamp recordings, slices were then transferred to a recording chamber and perfused with ascorbic-acid free ACSF at 28-30 °C.

### Electrophysiology

Voltage-clamp cell-attached and voltage/current-clamp whole-cell recordings were performed from LHb neurons in sagittal slices containing LHb using patch pipettes (3-6 MOhms) and a patch amplifier (MultiClamp 700B) under infrared-differential interference contrast microscopy. Data acquisition and analysis were carried out using DigiData 1440A, pCLAMP 10 (Molecular Devices). Signals were filtered at 3 kHz and digitized at 10 kHz. To assess LHb spontaneous activity and LHb neuronal excitability, cells were patch clamped with potassium gluconate-based internal solution (130 mM K-gluconate, 15 mM KCl, 4 mM adenosine triphosphate (ATP)-Na^+^, 0.3 mM guanosine triphosphate (GTP)-Na^+^, 1 mM EGTA, and 5 mM HEPES, pH 7.28, 275-280 mOsm) in slices perfused with ACSF. Spontaneous neuronal activity and action-potential (AP) firing patterns (tonic, bursting) were assessed in cell-attached recordings in voltage-clamp mode at V=0 for ∼2 min recording as previously described ^39, 52, 53^. LHb excitability experiments were performed either with intact fast-synaptic transmission to evaluate neuronal excitability or with the blockade of fast-synaptic transmission using DNQX (10µM), picrotoxin (100 µM), and D-APV (50 µM) in the ACSF to assess intrinsic excitability. LHb neurons were given increasingly depolarizing current steps at +10pA intervals ranging from +10pA to +100pA, allowing us to measure AP generation in response to membrane depolarization (5 sec duration). Current injections were separated by a 20s interstimulus interval and neurons were kept at ∼-65 to -70 mV with manual direct current injection between pulses. Resting membrane potential (RMP) was assessed immediately after achieving whole-cell patch configuration in current clamp mode. Input resistance (Rin) was measured during a -50pA step (5s duration) and calculated by dividing the steady-state voltage response by the current-pulse amplitude (-50pA) and presented as MOhms (MΩ). The number of APs induced by depolarization at each intensity was counted and averaged for each experimental group. As previously described^50^, AP number, AP threshold, fast and medium after-hyperpolarization amplitudes (fAHP and mAHP), AP halfwidth, AP amplitude were assessed using Clampfit and measured at the current step that was sufficient to generate the first AP/s. Hyperpolarization-activated cation currents (Ih) were evoked in LHb neurons that were voltage-clamped at -50mV by 2 sec voltage-steps of increasing amplitudes from −50 to −120 mV in steps of 10 mV. Under current-clamp potential recordings, sag potentials were elicited by 2 sec current-steps from -20 pA to -70pA in 10pA increases from a holding potential of -70mV. The amplitude of Ih current or sag potential was calculated as the difference of the peak and the steady state of the current or membrane potential induced in response to voltage/current steps, respectively.

To record M-currents, we used a standard deactivation protocol where LHb neurons were voltage-clamped at -60mV and received a 300 ms pre-pulse to -20 mV followed by 500 ms voltage steps from -30 to -75 mV in 5 mV increments. We calculated the amplitude of M-currents relaxation or deactivation as described before ^54^ by determining current relaxation which was the difference between the instantaneous (10 ms) and steady state (475 ms) of the current trace in response to voltage steps. The cell input resistance and series resistance were monitored through all the experiments and if these values changed by more than 10%, data were not included.

### Drugs

For all drug experiments, stock solutions for CRFR1 antagonists (antalarmin hydrochloride, Tcoris#2778 and NBI-35965 hydrochloride, Tocris#3100) were prepared in distilled water and diluted (1:1000) to final concentration in ACSF of 1 μM. LHb slices were incubated in the presence of vehicle/CRFR1 antagonists and also perfused in vehicle (ACSF)/CRFR1 antagonist-containing ACSF during drug-related excitability recordings.

### Statistics

Values are presented as mean ± SEM. The threshold for significance was set at *p < 0.05 for all analyses. All statistical analyses of data were performed using GraphPad Prism 10. Data from male and female mice were analyzed and reported separately to detect differences in sham versus mTBI. For detecting the difference between sham and mTBI mice in distribution of silent, tonic or bursting LHb neurons in spontoons activity and of escape and action-locking behaviors in looming shadow task, we used Chi-square tests. For depolarization-induced LHb excitability, I-V plot experiments and behavioral analysis of social interaction tests and elevated zero maze, two/three-way ANOVA were used. To detect the difference in intrinsic passive and active membrane properties and latencies to grooming in sucrose splash tests and escape/action-locking defensive behaviors in looming shadow task, we used two-tailed unpaired Student’s t-tests.

## Results

### mTBI increased LHb spontaneous activity and neuronal excitability in both male and female mice while inducing sexually dimorphic intrinsic plasticity

Previously, we have shown that mTBI results in persistent increases in spontaneous LHb tonic activity while decreasing LHb bursting in male mice^39^, although it was unclear how this model of mTBI affects LHb activity in female mice. Here, we evaluated the effects of mTBI on LHb spontaneous LHb activity (Figure 1A-B, D), LHb neuronal excitability (Figure 2A-B) and intrinsic excitability (Figure 2C-D) in LHb slices from sham and mTBI male and adult mice ∼4 weeks post-injury. mTBI increased the overall LHb spontaneous tonic activity while decreasing spontaneous LHb neuronal firing in bursting mode in cell-attached voltage-clamp recordings in male mice as we previously reported^39^, but also resulted in similar changes in female mice (Figure 1B, males: **p<0.01, Figure 1D, females: *p<0.05, Chi squared test). Consistently, LHb neurons of mTBI male and female mice also exhibited significantly higher neuronal excitability in intact synaptic transmission compared to those from sham mice although mTBI-induced LHb hyperexcitability was more pronounced in male mice (Figure 2 A, males: F (1, 836) = 63.46, ****p<0.0001; Figure 2B, females: F (1, 680) = 24.60, ****p<0.0001; sex effect in comparison between mTBI male group from Figure 2A and mTBI female group from Figure 2B: F (1, 806) = 63.69, ****p<0.0001, 2-way ANOVA). mTBI did not alter intrinsic membrane properties including RMP, Rin, fAHP, mAHP, AP amplitude, AP threshold and AP half-width (measurements extracted from intact excitability recordings in male or female mice, Supplemental Figures 1 and 2). Since neuronal excitability is dependent on both the synaptic inputs that LHb neurons receive and the intrinsic neuronal properties of LHb neurons, we then evaluated LHb intrinsic excitability in response to depolarization with blocked fast AMPAR, NMDAR and GABA_A_R-mediated transmission. We observed sex differences in the effects of mTBI where mTBI significantly increased LHb intrinsic excitability in male but not female mice (Figure 2 C, males: F (1, 203) = 22.51, ****p<0.0001; Figure 2D, females: F (1, 270) = 0.4947, p=0.4824, 2-way ANOVA). Interestingly, while mTBI did not alter any of the active and passive intrinsic membrane properties measured from intrinsic excitability recordings in male mice (Supplemental Figure 3), we detected a more depolarized voltage threshold for AP initiation coincident with higher levels of mAHPs in LHb neurons of mTBI female mice compared to those from sham female mice (Supplemental Figure 4, females: AP threshold, *p<0.05; mAHPs, **p<0.01, unpaired Student’s t test).

**Figure 1.**
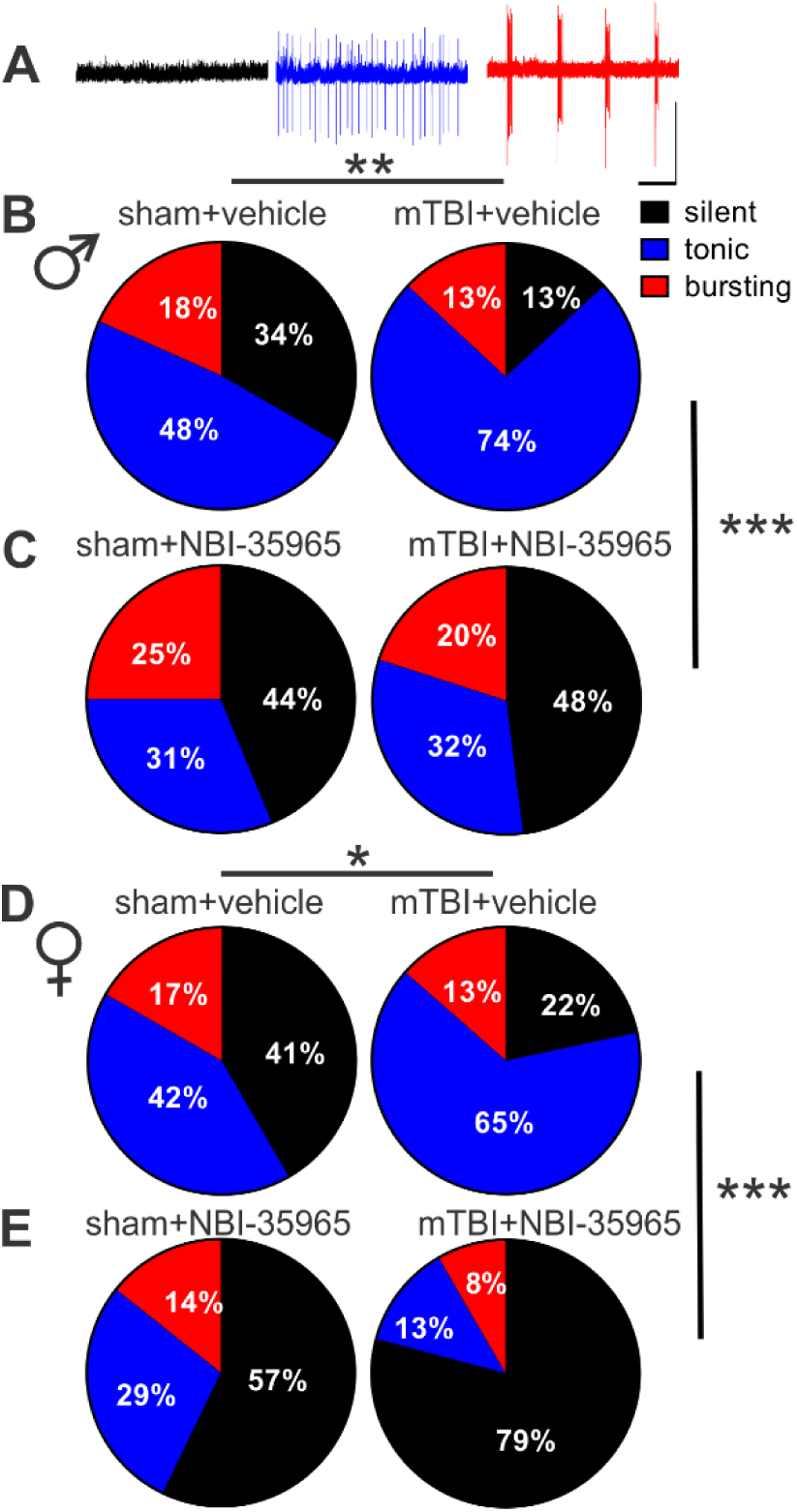
*In vitro* CRFR1 inhibition attenuated mTBI-induced increases in LHb tonic activity and decreases in LHb bursting in both male and female mice. Representative traces (**A**) and pie charts (**B-E**) of voltage-clamp cell-attached recordings (V=0 mV) of spontaneous neuronal activity with the percent distributions of silent (black), tonic (blue), or bursting (red) in LHb slices preincubated with either vehicle or NBI-35965 from male and female sham and mTBI mice (**B-C**, males: sham+vehicle, n=60/14; mTBI+vehicle, n=69/15, sham+NBI-35965, n=16/5, mTBI+NBI-35965, n=25/5; **D-E**, females: sham+vehicle, n=49/12; mTBI+vehicle, n=74/14, sham+NBI-35965, n=14/5, mTBI+NBI-35965, n=24/5). In this and subsequent electrophysiology graphs n represents the number of recorded cells/mice, *p<0.05, **p<0.01, ***p<0.001 by Chi squared tests.

**Figure 2:**
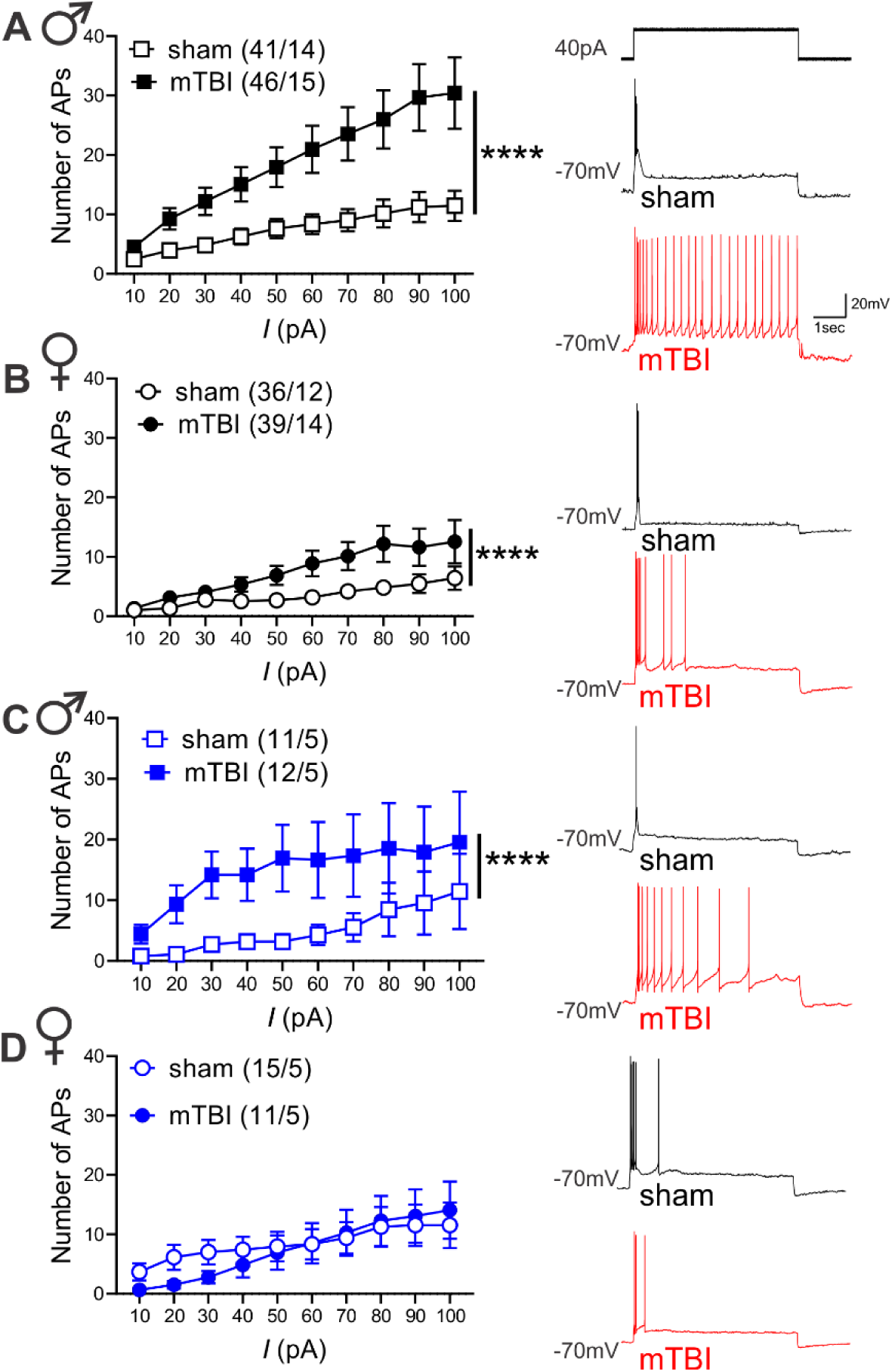
mTBI increased LHb neuronal excitability in male and female mice while only enhancing intrinsic excitability in male mice. (**A-B**) AP recordings in response to depolarizing current steps in LHb neurons in intact synaptic transmission and representative traces (sham: black, mTBI: red) from male/female sham and mTBI mice (male sham: black open square symbols, n=41/14, male mTBI: black filled square symbols, n=46/15, female sham: black open round symbols, 36/12, female mTBI: black filled round symbols, n=39/14). (**C-D**) AP recordings in response to depolarizing current steps in LHb neurons with fast synaptic transmission blocked and representative traces (sham: black, mTBI: red) from male/female sham and mTBI mice (male sham: blue open square symbols, n=11/5, male mTBI: blue filled square symbols, n=12/5, female sham: blue open round symbols, 15/5, female mTBI: blue filled round symbols, n=11/5); ****p<0.0001, 2-way ANOVA.

To further explore possible sexually dimorphic mechanisms underlying mTBI-induced intrinsic plasticity, we further evaluated the effects of mTBI on two membrane ionic currents (Ih currents and M-currents) that are abundantly expressed in LHb neurons and can potently regulate LHb activity where increases in Ih currents and decreases in M-currents are shown to promote LHb intrinsic excitability and LHb bursting ^27, 53, 55–59^. For this, we recorded Ih currents, sag potentials, hyperpolarization-induced rebound bursting and M-currents in response to hyperpolarizing and depolarizing current steps in LHb neurons of male and female sham and mTBI mice (Figures 3 and 4). Although LHb neurons from male and female mTBI mice displayed less spontaneous bursting activity as shown in Figure 1B and 1D, they fired more rebound bursts in response to hyperpolarization without any change in the amplitude of sag potentials (Figure 3B-C, males: rebound bursts, F (1, 397) = 47.44, ****p<0.0001, sag potentials, F (1, 406) = 0.4425, p=0.5063; Figure 4B-C, females: rebound bursts, F (1, 441) = 23.66, ****p<0.0001, sag potentials, F (1, 429) = 1.402, p=0.2370, 2-way ANOVA). Interestingly, mTBI enhanced Ih currents while slightly but significantly decreasing M-currents in LHb neurons of male mice (Figure 3A, D, Ih current: F (1, 434) = 8.188, **p<0.01, M-current: F (1, 520) = 3.955, *p<0.05, 2-way ANOVA). On the other hand, mTBI significantly enhanced M-currents in LHb neurons of female mice without affecting Ih currents (Figure 4D, Ih current: F (1, 516) = 0.2608, p=0.6098, M-current: F (1, 408) = 6.969, **p<0.01, 2-way ANOVA). Overall, our findings suggest that mTBI promotes LHb tonic hyperactivity, hyperexcitability and rebound bursting in male mice and female mice while inducing sex-dependent intrinsic plasticity.

**Figure 3.**
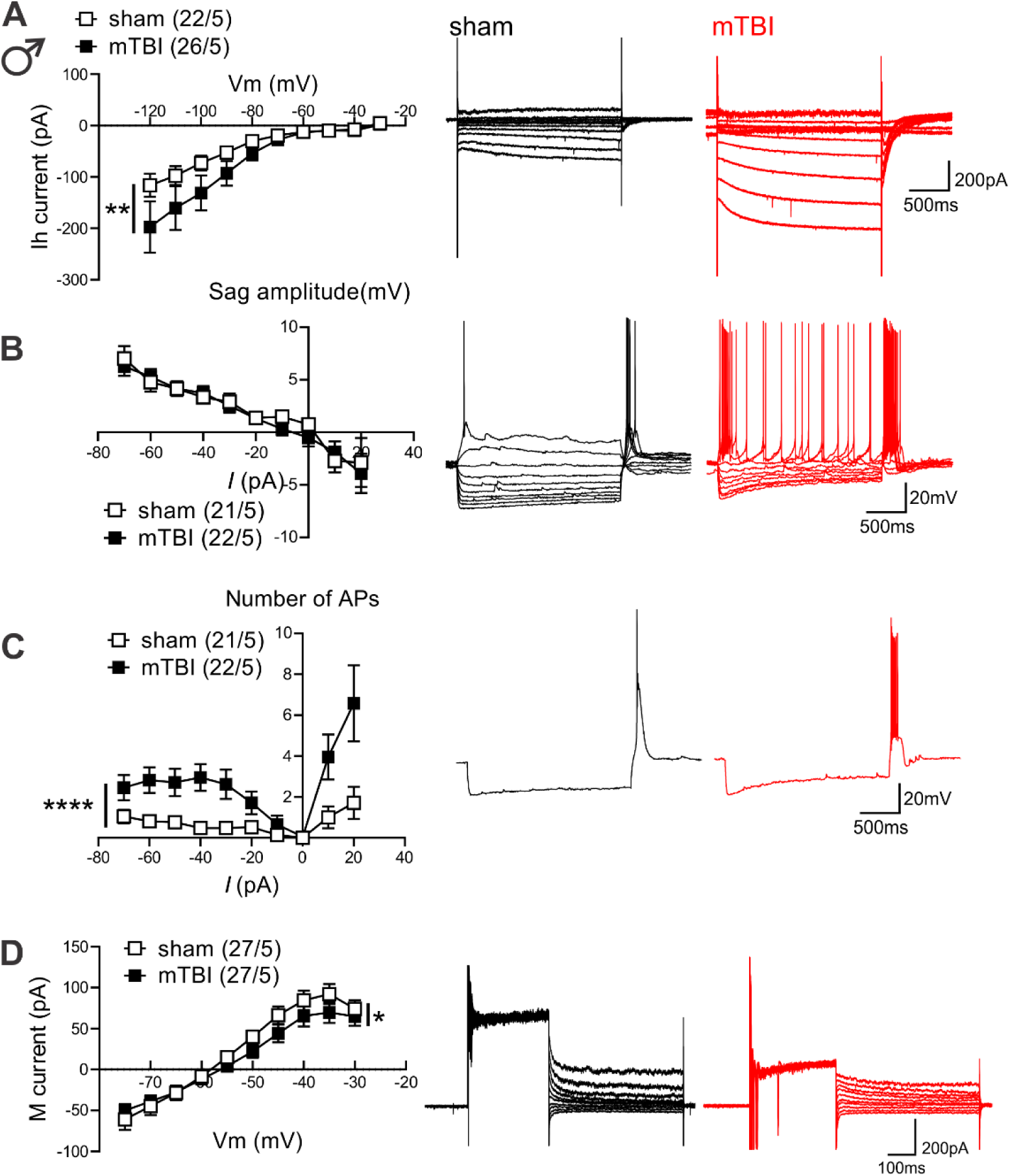
mTBI increased hyperpolarization-induced rebound bursts coincident with larger Ih currents and reduced M-currents in LHb neurons of male mice. **(A)** shows *I–V* relationship of Ih currents and representative traces (sham: black, mTBI: red) obtained from voltage-clamp recordings in LHb neurons from male sham (black open square symbols, n=22/5) and mTBI (black filled square symbols, n=26/5) mice. **(B)** shows *I–V* relationship of sag potentials and representative traces (sham: black, mTBI: red) obtained from current-clamp recordings in LHb neurons of male sham (black open square symbols, n=21/5) and mTBI (black filled square symbols, n=22/5) mice. **(C)** shows average number of APs in rebound bursts triggered in response to hyperpolarizing and depolarizing current steps in current-clamp recordings of sag potentials in **B** with representative traces of hyperpolarization-induced rebound bursts (sham: black, mTBI: red) in response to a -70pA hyperpolarizing current injection in LHb neurons of male sham (black open square symbols, n=21/5) and mTBI (black filled square symbols, n=22/5) mice. **(D)** shows *I–V* relationship of M-currents and representative traces (sham: black, mTBI: red) obtained from voltage-clamp recordings in LHb neurons from male sham (black open square symbols, n=22/5) and mTBI (black filled square symbols, n=26/5) mice; *p<0.05, **p<0.01, ****p<0.0001, 2-way ANOVA.

**Figure 4.**
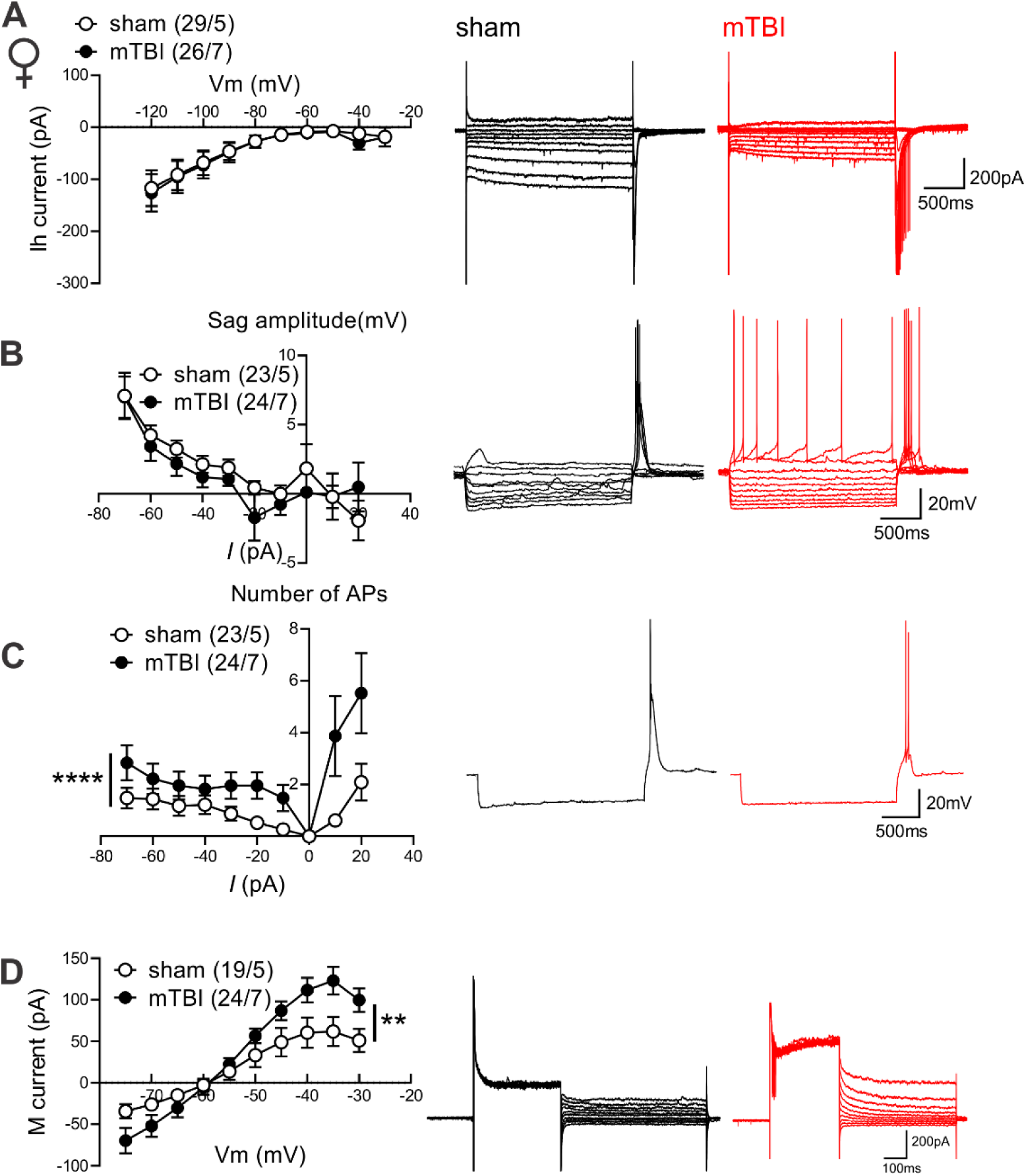
mTBI increased hyperpolarization-induced rebound bursts while enhancing M-currents in LHb neurons of female mice. **(A)** shows *I–V* relationship of Ih currents and representative traces (sham: black, mTBI: red) obtained from voltage-clamp recordings in LHb neurons from female sham (black open round symbols, n=29/5) and mTBI (black filled round symbols, n=26/7) mice. **(B)** shows *I– V* relationship of sag potentials and representative traces (sham: black, mTBI: red) obtained from current-clamp recordings in LHb neurons of female sham (black open round symbols, n=23/5) and mTBI (black filled round symbols, n=24/7) mice. **(C)** shows average number of APs in rebound bursts triggered in response to hyperpolarizing and depolarizing current steps in current-clamp recordings of sag potentials in **B** with representative traces of hyperpolarization-induced rebound bursts (sham: black, mTBI: red) in response to a -70pA hyperpolarizing current injection in LHb neurons of female sham (black open round symbols, n=23/5) and mTBI (black filled round symbols, n=24/7) mice. **(D)** shows *I–V* relationship of M-currents and representative traces (sham: black, mTBI: red) obtained from voltage-clamp recordings in LHb neurons from female sham (black open round symbols, n=19/5) and mTBI (black filled round symbols, n=24/7) mice; *p<0.05, **p<0.01, ****p<0.0001, 2-way ANOVA.

### mTBI-induced potentiation of intra-LHb CRF-CRFR1 signaling contributes to LHb hyperexcitability in male and female mice following mTBI

Previously, we demonstrated that CRF generally promotes LHb intrinsic excitability in rodent LHb ^50, 51^, however it was unclear whether mTBI engages endogenous CRF/CRFR1 signaling tone which could then contribute to alterations in the overall LHb neuronal activity in mTBI. To explore this possibility, we first pre-incubated LHb slices from sham and mTBI male mice with the selective CRFR1 antagonist (antalarmin, 1μM) and continued to bath-apply LHb slices with antalarmin which we previously showed to block the excitatory effects of CRF in the LHb^50^. We found that in the continuous presence of antalarmin, LHb tonic hyperactivity (Supplementary Figure 5A-B) and LHb hyperexcitability (Supplementary Figure 5C) in mTBI male mice became normalized to comparable levels observed in male sham mice (Supplementary Figure 5, A-B, effect of mTBI: **p<0.01, effect of antalarmin: ****p<0.0001, Chi squared tests; C, effect of mTBI: F (1, 600) = 11.02, ***p<0.001, effect of antalarmin: F (1, 600) = 17.22, ****p<0.0001, mTBI x antalarmin interaction: F (1, 600) = 14.79, ***p<0.001, three-way ANOVA). We then used a second selective CRFR1 antagonist, NBI-35965 (1μM), and LHb slices from male and female sham and mTBI mice were pre-incubated and perfused with this antagonist. Similar to antalarmin, NBI-35965 was also able to reverse the augmented LHb spontaneous tonic activity (Figure 1 B-E) and neuronal excitability (Figure 5) observed in male and female mTBI mice to control sham levels (Figure 1B-C, males: effect of mTBI: **p<0.01, effect of NBI-35965: ***p<0.001, Figure 1D-E, females: effect of mTBI: *p<0.05, effect of NBI-35965: ***p<0.001, Chi squared tests) (Figure 5A, males: effect of mTBI: F (1, 866) = 11.54, ***p<0.001, effect of NBI-35965: F (1, 866) = 21.70, ****p<0.0001, mTBI x NBI-35965 interaction: F (1, 866) = 36.49, ****p<0.0001; Figure 5B, females: effect of mTBI: F (1, 930) = 4.117, *p<0.05, effect of NBI-35965: F (1, 930) = 9.448, **p<0.01, mTBI x NBI-35965 interaction: F (1, 930) = 14.73, ***p<0.001, three-way ANOVA). Of note, we have combined all of the control data for neuronal excitability from the recordings of male sham mice interleaved with CRCR1 antagonist applications (either antalarmin or NBI-35965) and represented the combined data in Figure 2A.

**Figure 5:**
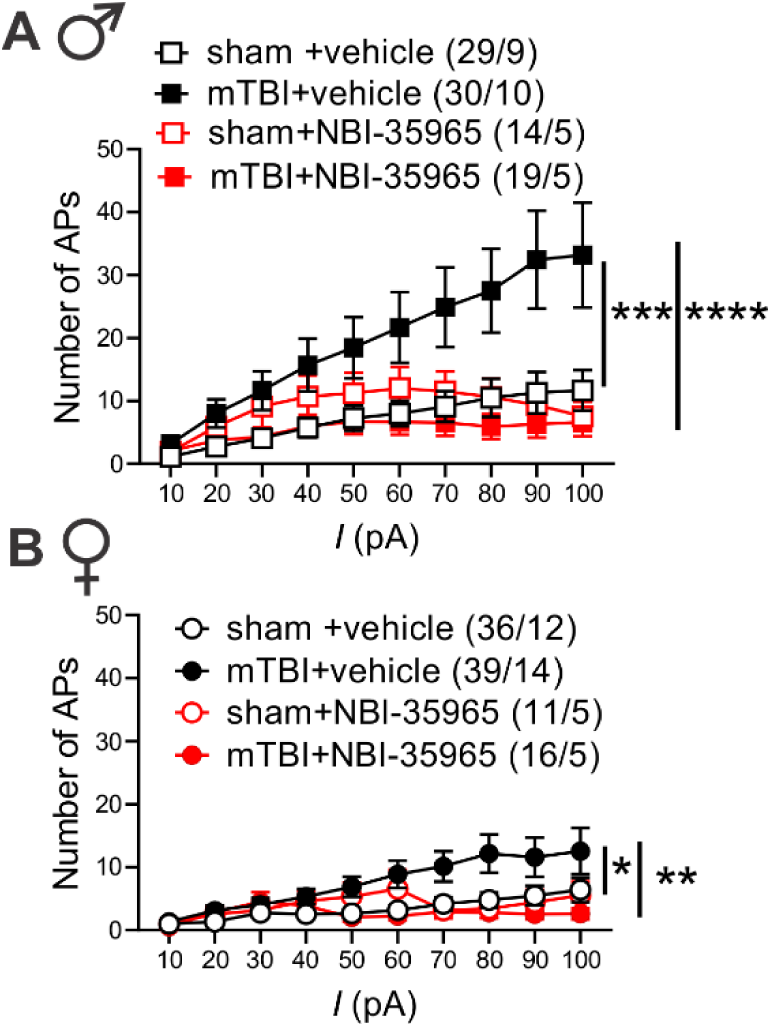
*In vitro* CRFR1 inhibition normalized mTBI-induced increases in LHb excitability in male and female mice. AP recordings in response to depolarizing current steps from LHb neurons in LHb slices of male (**A**) or female (**B**) sham and mTBI mice preincubated and perfused with either vehicle or NBI-35965 [**A,** males: sham+vehicle (black open square symbols, n=29/9); mTBI+vehicle (black filled square symbols, n=30/10) , sham+NBI-35965 (red open square symbols, n=14/5), mTBI+NBI-35965 (red filled square symbols, n=19/5); **B**, females: sham+vehicle (black open round symbols, n=36/12); mTBI+vehicle (black filled round symbols, n=39/14), sham+NBI-35965 (red open round symbols, n=11/5), mTBI+NBI-35965 (red filled round symbols, n=16/5); *p<0.05, **p<0.01, ***p<0.001, ****p<0.0001, 3-way ANOVA.

### mTBI induced deficits in motivation to self-care grooming and defensive behaviors in both male and female mice but only triggered social deficits in male mice

To establish a complete readout of depressive-, anxiety-, and PTSD-like behaviors in this model of mTBI, we performed behavioral assays, the sucrose splash test (only in female mice, Figure 7A-B), the social interaction test (Figure 6A-B and Figure 7C-D), the elevated zero maze (Figure 6C and Figure 7E) and the looming-induced innate defensive behaviors (Figure 6D-E and Figure 7F-G), in male and female sham and mTBI mice. Similar to what we previously observed in male mTBI mice^39^, mTBI female mice exhibited increased latency to grooming in sucrose splash test without any alteration in their total grooming behavior, suggesting that this mTBI model promotes similar motivational deficits in self-care grooming in male and female mice (Figure 7A-B, females, latency: t=4.095, df=17, ***p<0.001, un-paired Student’s t-tests). We also replicated our previous findings^40^ that this model of mTBI persistently decreases social interaction with a same-sex conspecific stranger in male mice. While male sham mice showed significant preference for exploring a male novel mouse and spent more time with a novel conspecific (social zone) over the non-social zone containing an empty cup (empty), indicative of substantial degree of sociability in male sham mice, male mTBI mice did not show this preference and spent a similar proportion of time in social and empty zones. Subsequently, male mTBI mice spent a significantly lower proportion of time in the social zone compared to male sham mice (Figure 6A, sociability: F (1, 36) = 6.425, *p<0.05, mTBI: F (1, 36) = 1.337, p=0.2553, sociability x mTBI interaction: F (1, 36) = 11.80, **p<0.01, two-way ANOVA). We also tested social novelty following mTBI (not tested before in our earlier study) in which we exposed sham and mTBI mice to a second stranger mouse that was placed in the empty zone (as a novel mouse) while the first stranger remained in the same social zone at the end of first session (as a familiar mouse). We found that male sham mice spent similar time exploring familiar or novel mice while male mTBI mice increased their interaction with the novel mouse (Figure 6B, novelty: F (1, 36) = 6.003, *p<0.05, mTBI: F (1, 36) = 0.3436, p=0.5614, novelty x mTBI interaction: F (1, 36) = 2.864, p=0.0992, two-way ANOVA). Female sham and mTBI mice spent similar time exploring social (the first conspecific stranger) and non-social (empty) zones in the first session of the social interaction test (Figure 7C, sociability: F (1, 36) = 2.730, p=0.1072, mTBI: F (1, 36) = 0.2229, p=0.6397, sociability x mTBI interaction: F (1, 36) = 0.9074, p=0.3472). On the other hand, in the second session female mTBI mice also spent more time exploring the novel mouse, avoiding the familiar mouse as we observed in male mTBI mice (Figure 7D, novelty: F (1, 35) = 10.66, **p<0.01, mTBI: F (1, 35) = 0.2431, p= 0.6250, novelty x mTBI interaction: F (1, 35) = 0.7998, p=0.3773, two-way ANOVA). To further test novelty seeking/risk-taking/anxiety-like behaviors, we performed elevated zero maze in which we observed that both male and female sham and mTBI mice significantly increased their time spent in closed versus open arms with no significant differences in time spent in closed versus open arms between sham and mTBI groups of male or female mice in this test (Figure 6C, arm effect: F (1, 56) = 394.7, ****p<0.0001, mTBI: F (1, 56) = 0.2877, p=0.5938, arm x mTBI interaction: F (1, 56) = 0.4434, p=0.5082; Figure 7E, arm effect: F (1, 36) = 467.2, ****p<0.0001, mTBI: F (1, 36) = 0.0004, p=0.9838, arm x mTBI interaction: F (1, 36) = 1.733, p=0.1964, two-way ANOVA).

**Figure 6.**
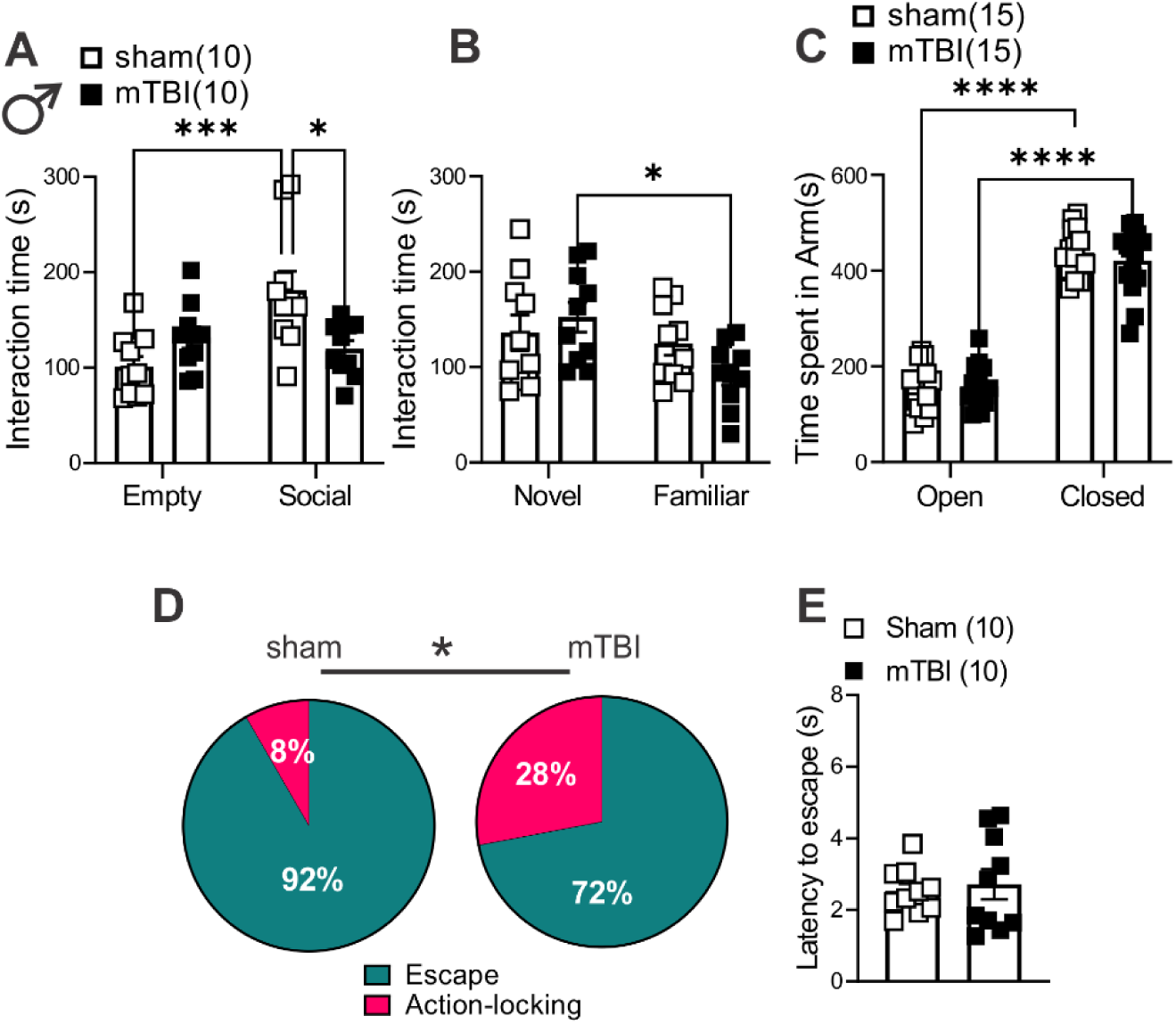
mTBI induced behavioral deficits in self-care grooming, social interaction and threat-evoked defensive behaviors in male mice. (**A**) shows time spent interacting with a novel conspecific (first stranger mouse, Social zone) or a non-social zone containing an empty cup (Empty) in three chamber social interaction tests in male sham (black open square symbols, n=10) and mTBI (black filled square symbols, n=10) mice. (**B**) shows time spent interacting with the first stranger mouse (Familiar) or a second stranger mouse (Novel) in three chamber social interaction tests in male sham (black open square symbols, n=10) and mTBI (black filled square symbols, n=10) mice. (**C**) shows time spent in open arms and closed arms in an elevated zero maze in male sham (black open square symbols, n=15) and mTBI (black filled square symbols, n=15) mice. (**D**) shows the percent distributions of escape (teal blue) or action-locking (magenta) behaviors in response to a looming shadow threat in male sham (n=10) and mTBI (n=10) mice. (**E**) shows latencies to escape responses to a looming shadow threat in male sham (black open square symbols, n=10) and mTBI (black filled square symbols, n=10) mice; * p<0.05, ***p<0.001, ****p<0.0001, 2-way ANOVA; Chi squared tests; * p<0.05.

**Figure 7.**
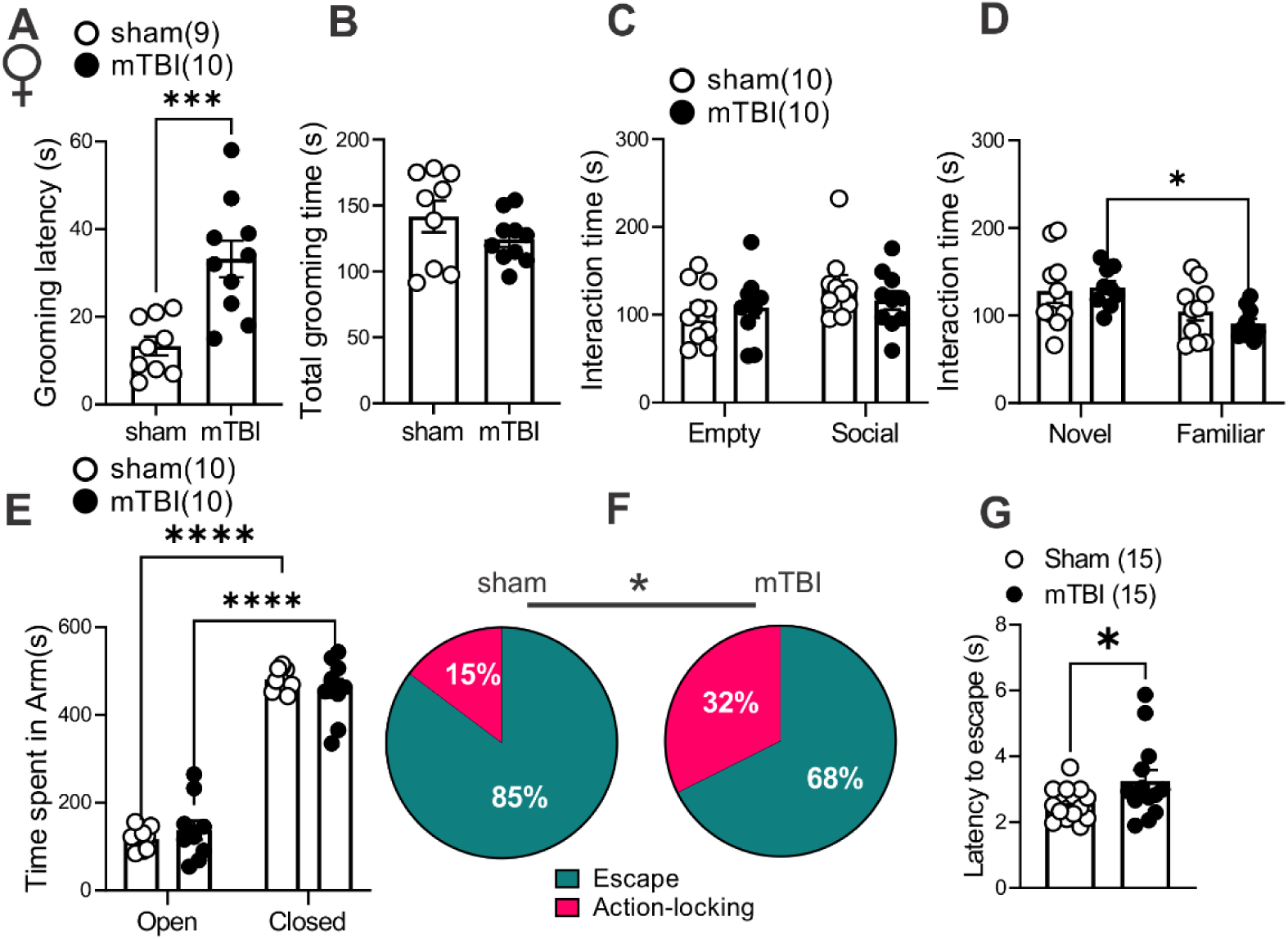
mTBI induced behavioral deficits in self-care grooming and threat-evoked defensive behaviors in female mice. (**A-B**) shows latency to grooming and total grooming time in sucrose splash tests in female sham (black open round symbols, n=9) and mTBI (black filled square symbols, n=10) mice. (**C**) shows time spent interacting with a novel conspecific (first stranger mouse, Social zone) or a non-social zone containing an empty cup (Empty) in three chamber social interaction tests in female sham (black open round symbols, n=10) and mTBI (black filled round symbols, n=10) mice. (**D**) shows time spent interacting with the first stranger mouse (Familiar) or a second stranger mouse (Novel) in three chamber social interaction tests in female sham (black open round symbols, n=10) and mTBI (black filled round symbols, n=10) mice. (**E**) shows time spent in open arms and closed arms in an elevated zero maze in female sham (black open round symbols, n=10) and mTBI (black filled round symbols, n=10) mice. (**F**) shows the percent distributions of escape (teal blue) or action-locking (magenta) behaviors in response to a looming shadow threat in female sham (n=15) and mTBI (n=15) mice. (**G**) shows latencies to escape responses to a looming shadow threat in female sham (black open round symbols, n=15) and mTBI (black filled round symbols, n=15) mice; * p<0.05, ***p<0.001, ****p<0.0001, 2-way ANOVA; Chi squared and Student’s t-tests; * p<0.05.

We also chose to pursue mouse defensive behaviors in the looming shadow test as a complex and ethologically relevant behavior that is evolutionarily conserved and with high translational value^60^. In this behavioral assay, mice adopt innate defensive behaviors to looming shadows approaching from above by active (escape-to-nest, the predominant behavior) or passive (action-locking, also referred as immobile-like or freezing-in-place when there is less confidence of escape to a safe shelter) behaviors. Therefore, we tested whether mTBI biases threat responses towards one of the defensive behaviors (increased action-locking or escape) and/or alters the latency to threat responses. We found that mTBI significantly increased action-locking responses in both male and female mice (Figure 6D and 7F, effect of mTBI: *p<0.05, Chi squared tests). mTBI also prolonged the latency to escape defensive behaviors in female but not male mice (Figure 6E and 7G, effect of mTBI in females: t=2.024, df=26, *p<0.05, Un-paired Student t-test). Overall, mTBI impaired normal defensive behaviors resulting in persistence of action-locking behaviors in both male and female mice and delayed defensive stress reactions only in females.

## Discussion

Previous works from our laboratories have established that LHb hyperactivity plays a causal role in motivational deficits in self-care behaviors in a repetitive closed head mTBI model^39^ as well as this model induces social anhedonia ^40^ in male mice suggesting that this preclinical model of mTBI may predict some aspects of mTBI-induced reward/motivational circuit dysfunction underlying long-term negative mood outcomes related to mTBI in humans. Here, we further characterized the neurobehavioral effects of this model in both young male and female mice with the time of injury during the transition from late adolescence to young adulthood (at ∼P56). We have focused on the exposure to mTBI injury at this critical age in mice given that older adolescent humans (age 15 to 19) represent one of the most at-risk populations for mTBI^1^. Our study uncovers that this model mostly induces similar neurophysiological and mood-related behavioral alterations in male and female mice including an overall LHb hyperactivity and hyperexcitability partly due to hypertrophy of intra-LHb CRF-CRFR1 signaling and intrinsic plasticity, as well as deficits at the level of the motivation to self-care behavior and approach/avoidance in social- and threat-evoked behavioral responses in male and female mice.

This model of mTBI resulted in increased LHb tonic activity while diminishing LHb bursting in female mice similar to what we previously reported in male mice ^39^. Consistent with the overall increased LHb tonic activity, we found that mTBI promoted LHb hyperexcitability in male and female mice. Furthermore, we observed that *in vitro* inhibition of CRFR1 within the LHb normalized mTBI-induced enhancement of both LHb spontaneous tonic activity and hyperexcitability in both sexes, suggesting that augmentation of CRF-CRFR1-mediated signaling within the LHb may provide a neuromodulatory mechanism by which mTBI triggers LHb hyperactivity in both male and female mice. CRF is widely known for its participation in stress-induced activation of the HPA axis, but CRF also acts within the brain where it directly regulates both positive and negative reinforcement of motivated behaviors in response to stressors through its receptors, CRF receptor types 1 and 2 (CRFR1, CRFR2) ^61–64^. This extra-hypothalamic CRF system includes brain regions whose activities are implicated in processing of reward and aversive information as well as in mental disorders and drug addiction ^65–68^. Up to 30% of TBI patients experience neuroendocrine dysfunction ^41–44^ with a high incidence of HPA axis dysregulation ^42,44^. Hypothalamic and extra-hypothalamic CRF systems are also responsive to mTBI injuries and have significant impact on stress-related neuronal responses and affective states following mTBI ^45–49^. The mouse LHb shows immunoreactivity and mRNA expression for both CRF and CRFR1 ^69, 70^, and receives inputs from the structures that contain CRF neurons including the paraventricular nucleus of hypothalamus (PVN) and the extrahypothalamic CRF system such as the bed nucleus of stria terminalis (BNST)^71^. Therefore, it will be worthwhile to examine whether CRF projections to the LHb are susceptible to modulation by mTBI injury and causally contribute to overall LHb hyperactivity in mTBI mice of both sexes.

Interestingly, CRF can also increase intrinsic excitability in both rat and mouse LHb, although the mechanisms underlying the excitatory effects of CRF on LHb intrinsic excitability may differ between species due to suppression of different types of potassium channels contributing to mAHPs in LHb neurons, i.e., SK channels in rat LHb^50^ versus M channels in mouse LHb^51^. Voltage gated K^+^ channel 7 (Kv7), also known as M-currents, have a relatively low activation voltage and rapidly open in response to small depolarizations near resting membrane potential, contributing to mAHPs and stabilization of neuronal membrane potential, thereby limiting excessive neuronal excitability^72, 73^. Activation of LHb M channels is also shown to decrease LHb neuronal activity and the anxiety-like phenotype induce by alcohol withdrawal in rats^59^. We have also shown a critical role for the scaffolding A-kinase anchoring protein (AKAP150) in regulation of synaptic transmission, plasticity and intrinsic excitability of LHb neurons^51^. We demonstrated that genetic disruption of AKAP150-anchored PKA increases LHb intrinsic excitability, and saturates and occludes the excitatory actions of CRF on intrinsic excitability in mouse LHb possibly through AKAP150-PKC dependent suppression of M-channels^74^. Multiple synaptic inputs to the LHb co-release both glutamate and GABA^75^, and CRF also reduces presynaptic glutamatergic and GABAergic transmission in mouse LHb which is consistent with the prior observations that CB1 receptors are expressed on both presynaptic glutamate and GABA terminals in the LHb^51, 76^. Therefore, CB1 receptor activation by eCBs downstream from augmented CRF signaling^50^ could decrease the probability of presynaptic glutamate and GABA release onto LHb neurons at distinct synaptic inputs to the LHb (e.g., lateral preoptic area) where the eCB-CB1 receptor-mediated suppression of presynaptic GABA release is assumed to be predominantly larger ^77^. Human studies of polymorphisms of *AKAP5* gene also suggests that AKAP5 plays an essential regulatory role in mood, cognitive control of anger, aggression and impulsivity ^78–81^. Therefore, in the context of mTBI-related mood deficits, it will be interesting to investigate whether mTBI-induced augmentation of CRF-CRFR1 signaling and LHb hyperactivity involve any dysregulation of AKAP150-mediated synaptic or CRF neuromodulatory functions within the LHb and its circuits that are known to regulate impulsive, depressive-, aversive- and drug-related behaviors.

While *in vitro* CRFR1 inhibition within the LHb neurons was effective in reversing mTBI-induced LHb hyperexcitability in both male and female mice, the LHb hyperexcitability (recorded with intact synaptic transmission) was notably more pronounced in male mTBI mice than female mTBI mice suggesting the possibility of an induction of sexually dimorphic intrinsic plasticity by mTBI. Consistent with this, we only observed intrinsic hyperexcitability (with synaptic transmission blocked) in the LHb neurons of male but not female mice. While we did not observe any significant change in mAHP measurements from intrinsic excitability of LHb neurons of male mTBI compared to sham mice, we detected a slight but significant reduction in M-currents induced by mTBI providing a molecular mechanism for induction of this intrinsic plasticity by mTBI in male mice. Interestingly, mTBI induced some changes in intrinsic properties of LHb neurons of female mice that could explain the lack of mTBI-induced intrinsic hyperexcitability and the smaller extent of intact LHb hyperexcitability in females. These include a more depolarized AP threshold and higher levels of mAHPs associated with significant potentiation of M-currents in LHb neurons of mTBI female mice compared to those from sham female mice. Enhancement of M-currents in LHb neurons may be a compensatory mechanism triggered in response to mTBI injury in female mice to counteract an exaggerated LHb hyperexcitability that was observed only in male mouse LHb following mTBI. In fact, mTBI also enhanced Ih currents in male but not female mice which could also contribute to LHb hyperexcitability by Ih-mediated depolarization and diminishing of GABA_A_-receptor mediated synaptic inhibition onto LHb neurons^82^. Almost all LHb neurons projecting to the VTA or raphe nuclei appear to express all four HCN subunits ^55^, however, a later study found that only a subset of glutamatergic medial LHb neurons with functional Ih are excited by dopamine-dopamine D4 receptor activation of Ih currents and these neurons project to the RMTg but not to the VTA ^56^. Therefore, there is a possibility of sexual dimorphism in differential circuit effects of mTBI on LHb projections to RMTg, VTA and raphe nuclei and LHb-mediated behaviors. Interestingly, both male and female mice exhibited an increase in hyperpolarization-induced rebound bursts. Mechanistically, increased Ih currents in LHb neurons of male mice can also promote LHb bursting by NMDA receptor activation that is required for bursting while enhanced potentiation of M-current may also provide the necessary hyperpolarization for activation of low-voltage-sensitive T-type calcium channels that also promote LHb bursting. Photo-stimulation of the optogenetic chloride pump, halorhodopsin, in LHb neurons triggers hyperpolarization-induced rebound bursts in the LHb which also promotes behavioral aversion and depression-like phenotype in the real-time place aversion assay in mice^27^. Therefore, we assume that under conditions in which GABAergic inhibition of LHb neurons occurs, heightened rebound bursting activity of LHb neurons in mTBI mice can drive negative affective states and depressive-like behaviors in mTBI male and female mice.

From behavioral standpoint, we found that this model of mTBI also diminishes motivation for self-care grooming in female mice, as we had observed in male mice^39^. Moreover, we replicated our previous finding that mTBI decreases social preference in male mice although this effect of mTBI was not observed in female mice. Of note, in the three chamber social interaction tests female sham mice also did not significantly increase their interaction with a conspecific stranger versus empty cup indicating that mTBI-induced social deficits in female mice could not reliably be excluded using this test. LHb plays an important role in social behaviors and conspecific interactions^83, 84^. For example, chemogenetic activation of LHb neurons is shown to diminish social interaction, and similarly optogenetic stimulation of mPFC-LHb pathway suppresses social preference^83^. Our model of mTBI indeed affects the anterior cingulate cortex which is considered the dorsal component of mPFC ^85^ and results in low levels of axonal damage in mPFC in this mTBI model^40^. This raises the possibility of the involvement of a dysregulated mPFC-LHb pathway in decreased social preference in male mice following mTBI that may indicate that male mice are more susceptible to this type of injury than female mice. Indeed, estrogen receptors are highly expressed in the LHb and estradiol can suppress LHb spontaneous neuronal activity^86^. A subpopulation of GABAergic interneurons is also identified in the LHb that are estrogen-receptive, can locally inhibit LHb glutamatergic neurons and regulate motivated behavior^87, 88^. Therefore, it is possible that compensatory mechanisms triggered by mTBI in female mice such as upregulation of M-currents in LHb neurons as well as estrogen and/progesterone-mediated neuroprotection in young female adult mice provides some level of resilience within social-relevant LHb pathways for females to prevent social interaction deficits. Curiously, while both male and female mTBI mice did not show any anxiety/risk-taking/novelty seeking behavior in the elevated zero maze tests, they exhibited an increased tendency to explore a novel mouse in the social novelty test. The high novelty seeking behavior is suggested to predict aggressive behavior where increased intrusion-evoked aggressive behaviors in male rats with heightened novelty seeking were observed with coincident diminished intrusion-induced c-fos activation in selected raphe serotonergic neurons^89^. It is not apparent whether the increased exploration of the second stranger mouse (the novel mouse) versus the first stranger mouse (the familiar mouse) in the social novelty test in mTBI mice indicates any types of pathological aggressive behaviors or territorial aggression. Of interest, activation of LHb glutamatergic projections to the dorsal raphe non-serotonergic neurons promotes exaggerated inter-male aggressive behaviors in mice as a result of social instigation when mice were pre-exposed to a rival male mouse^90^. Although, preclinical models of female aggression are scarce, it is possible that mTBI female mice that exhibited depressive phenotype in the sucrose splash test also show pathological aggression in behavioral tests of aggression of social instigation due to LHb hyperactivity and the possible suppression of the raphe serotonergic system. Thus, it will be worthwhile to tease apart the contribution of LHb-raphe pathway dysregulation in distinct aspects of social novelty seeking and socially primed-aggressive behaviors in male and female mice following mTBI.

We also chose to evaluate mouse defensive behaviors in the looming shadow test after mTBI as a complex and ethologically relevant behavior that is evolutionarily conserved and with high translational value^60^. The looming shadow task offers face validity as similar threat responses of active fleeing (flight) or passive staying (freezing) strategies exist in humans in response to imminent threats, and the aberrant rigidity in defensive strategies such as persistence of either freezing or escape behaviors and prolonged threat response reactivities are observed in stress-related psychopathologies including anxiety and PTSD^91–96^ ; common comorbidities associated with mTBI. The use of predator threats has also been proven more valuable for generating and evaluating PTSD-like phenotypes^97^. We observed that mTBI impaired defensive behaviors in the looming shadow task by shifting the innate defensive behaviors towards more passive action-locking rather than escape behaviors in response to an aerial threat, in both male and female mice, and also resulted in prolonged latencies to escape responses in female mice. Aberrant threat responses and defensive behaviors, specifically increased in passive (freezing) but not active (escape) defensive behaviors are attributed to stress-related psychiatric morbidity risk after trauma including depression, anxiety and PTSD^92, 98–100^ although high trait anxious individuals also exhibit more attentional biases towards threat reflecting abnormal reactivity to threat cues and hypervigilance that may promote escape behavior^101, 102^. Prolonged latency to defensive behaviors in looming shadow task has also been observed following an early life adversity model as a risk for psychiatric disorders^103^. Importantly, during threat-provoked defensive responses the LHb becomes engaged. For example, exposure of mice to a predator or predator odor increases the expression of the immediate early gene c-Fos in the LHb suggesting threat-evoked increased LHb neuronal activity ^87, 104^. Additionally, optogenetic activation of LHb glutamatergic terminals to laterodorsal tegmentum GABAergic interneurons promotes fear-like responses (freezing) behaviors in response to predator odor in mice ^104^. Dynamic activity of LHb neuronal populations in behaving mice exposed to the looming shadow has also revealed time-locked excitation and inhibition responses of distinct clusters of LHb neurons with escape and action-locked behaviors, respectively^105^. Altogether, we provided several lines of evidence for translational validity of a preclinical model of mTBI in male and female mice that is associated with LHb hyperactivity and intra-LHb CRF dysregulation, associated with a lack of motivation in self-care, disrupted social behavior and aberrant threat responses that are core symptoms of many psychiatric conditions including depression, anxiety and PTSD. Our model enables for future investigations into mTBI-induced maladaptive changes in molecular, synaptic and neuronal mechanisms at the level of distinct LHb circuits with implications for psychiatric disorders in mTBI.

## Acknowledgments

The opinions and assertions contained herein are the private opinions of the authors and are not to be construed as official or reflecting the views of the Uniformed Services University of the Health Sciences or the Department of Defense or the Government of the United States. Behavioral testing and analysis were performed in the Preclinical Behavior and Modeling Core at the Uniformed Services University.

## Authorship contribution

FN and WF were responsible for the study concept and design. RA contributed to the experimental design for the mTBI injury. MT and WF established the looming shadow task apparatus in Nugent laboratory. WF, SS, ET, SG, MT and FN contributed to the acquisition of animal data. FN, WF, ET, SS, SG, MT, JW, BC and AS assisted with data analysis and interpretation of findings. WF and FN wrote the initial draft of the manuscript. All authors critically reviewed the content and approved final version of manuscript for submission. The authors acknowledge Dr. Yeonho Kim, Dr. Amanda Fu and Laura Tucker at the USU Preclinical Modeling and Behavior Core for supporting the implementation of sham and mTBI procedures and behavioral studies.

## Data Sharing

The data that support the findings of this study are available on request from the corresponding author.

