## Supplemental Figures for "Effects of Repetitive Mild Traumatic Brain Injury on Corticotropin-Releasing Factor Modulation of Lateral Habenula Excitability and Motivated Behavior"

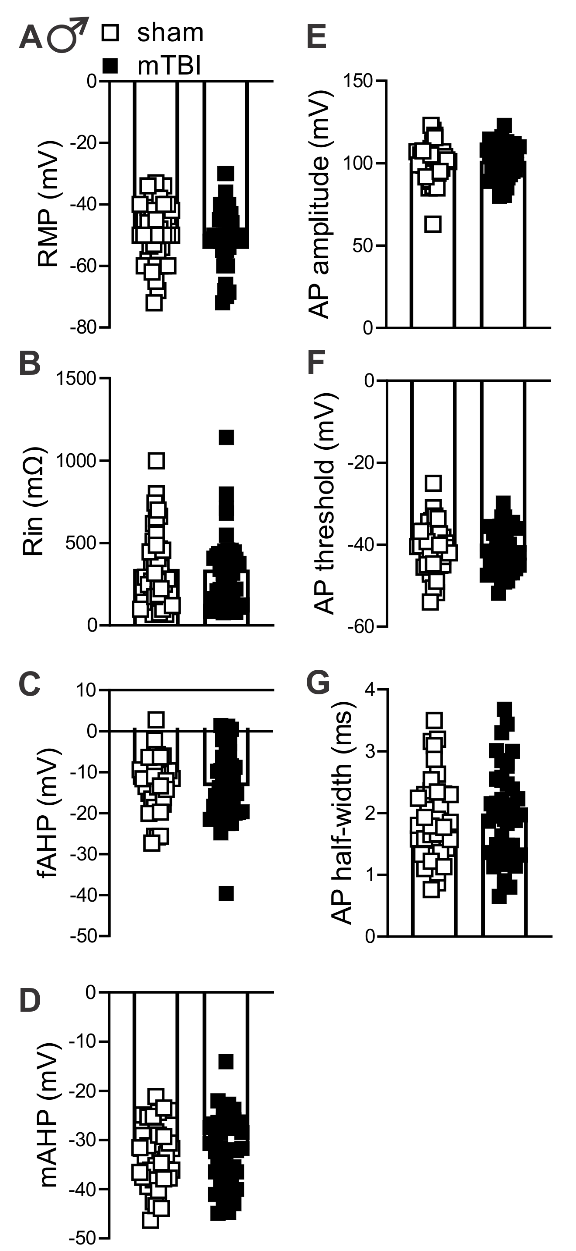

**Supplemental Figure 1. mTBI did not affect intrinsic membrane properties of LHb neurons in intact synaptic transmission in male mice.** (**A-G**) measurements of RMP, Rin, fAHP, mAHP, AP amplitude, AP threshold, and AP half width in LHb neurons derived from AP recordings in response to depolarization in Figure 2A from male sham (black open square symbols, n=36/12) and mTBI (black filled square symbols, n=39/14) mice.

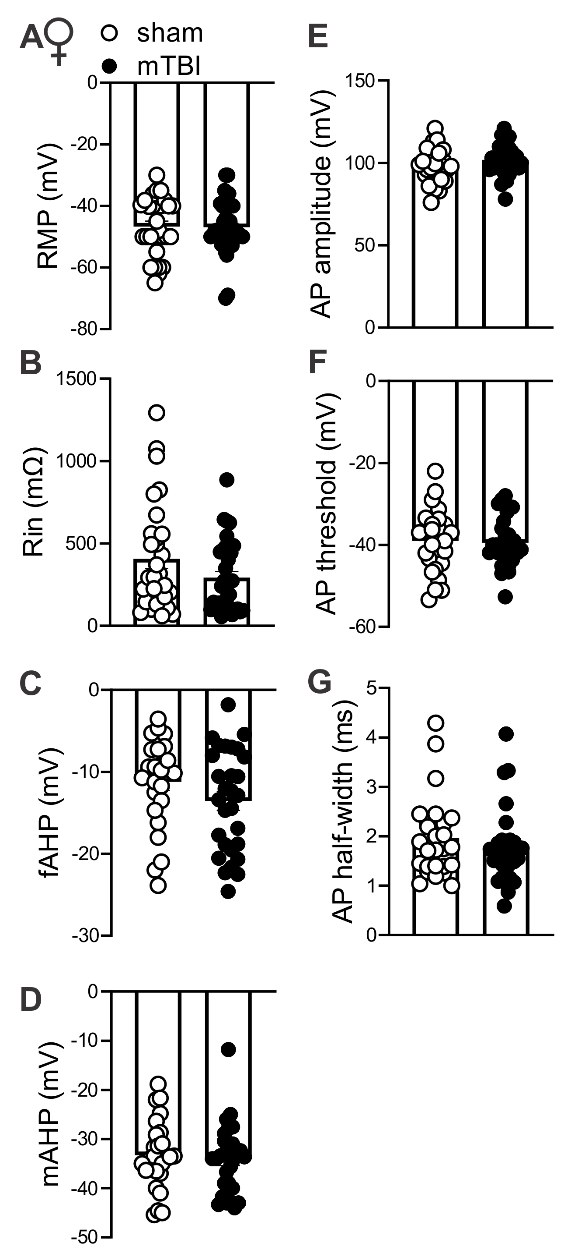

**Supplemental Figure 2. mTBI did not affect intrinsic membrane properties of LHb neurons in intact synaptic transmission in female mice.** (**A-G**) measurements of RMP, Rin, fAHP, mAHP, AP amplitude, AP threshold, and AP half width in LHb neurons derived from AP recordings in response to depolarization in Figure 2B from female sham (black open round symbols, n=36/12) and mTBI (black filled round symbols, n=39/14) mice.

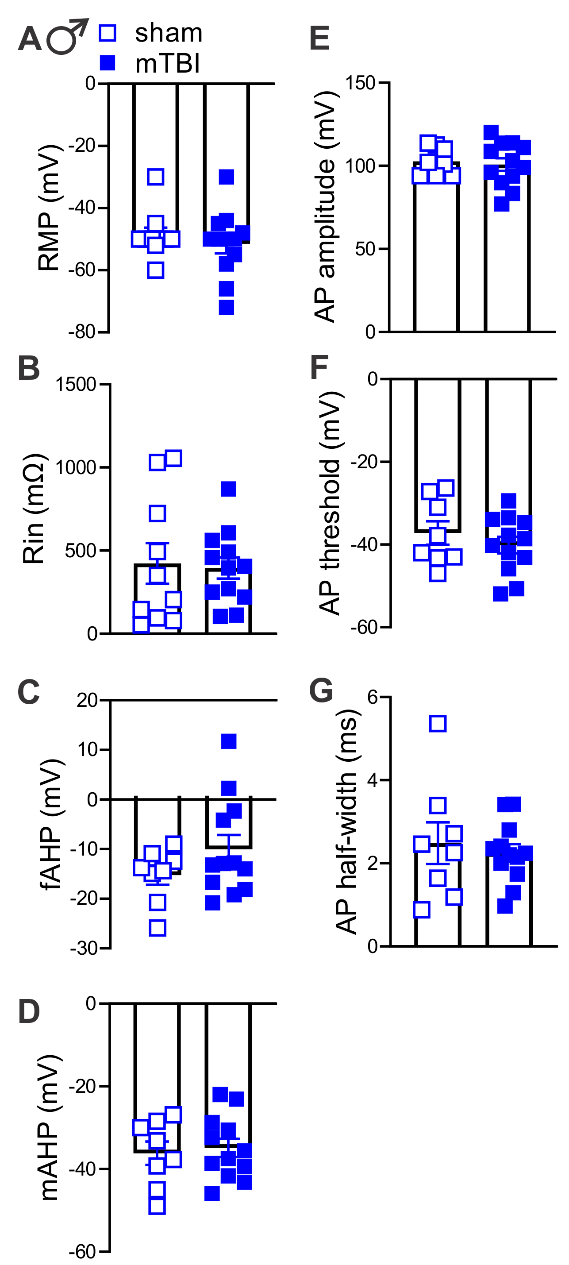

**Supplemental Figure 3. mTBI did not affect intrinsic membrane properties of LHb neurons with fast synaptic transmission blocked in male mice.** (**A-G**) measurements of RMP, Rin, fAHP, mAHP, AP amplitude, AP threshold, and AP half width in LHb neurons derived from AP recordings in response to depolarization in Figure 2C from male sham (black open square symbols, n=11/5) and mTBI (black filled square symbols, n=12/5) mice.

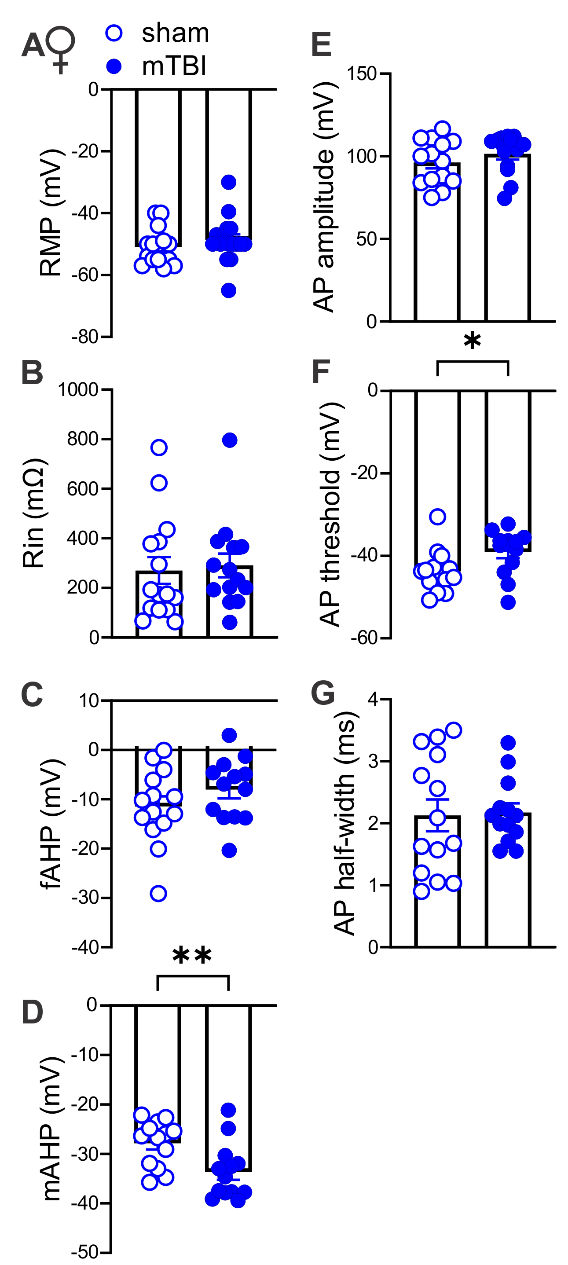

**Supplemental Figure 4. mTBI increased mAHPs and resulted in a more depolarized AP threshold in LHb neurons with fast synaptic transmission blocked in female mice.** (**A-G**) measurements of RMP, Rin, fAHP, mAHP, AP amplitude, AP threshold, and AP half width in LHb neurons derived from AP recordings in response to depolarization in Figure 2D from female sham (blue open round symbols, n=15/5) and mTBI (blue filled round symbols, n=11/5) mice; *p<0.05, **p<0.01, unpaired Student’s t-tests.

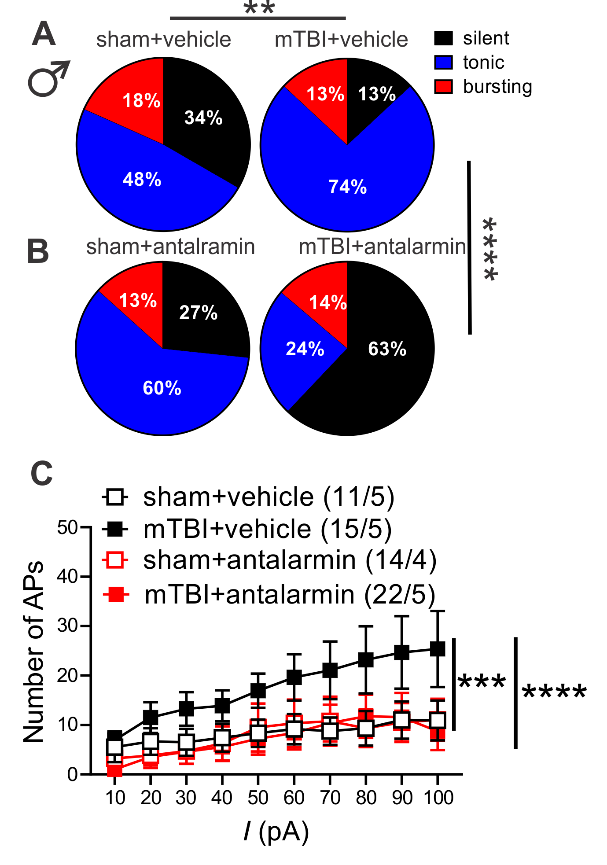

**Supplemental Figure 5. *In vitro* CRFR1 inhibition by antalarmin normalized mTBI-induced increases in LHb spontaneous tonic activity and excitability in male mice.** (**A-B**) and pie charts of voltage-clamp cell-attached recordings (V=0 mV) of spontaneous neuronal activity with the percent distributions of silent (black), tonic (blue), or bursting (red) in LHb slices preincubated with either vehicle or antalarmin from male sham and mTBI mice. Note that data from sham+vehicle and mTBI+vehicle groups in this graph are identical with those control groups represented in Figure 1B for NBI-35965 experiments (males: sham+vehicle, n=60/14; mTBI+vehicle, n=69/15, sham+antalarmin, n=15/4, mTBI+NBI-35965, n=29/5). (**C**) AP recordings in response to depolarizing current steps from LHb neurons in LHb slices of male sham and mTBI mice preincubated and perfused with either vehicle or antalarmin [sham+vehicle (black open square symbols, n=11/5); mTBI+vehicle (black filled square symbols, n=15/5) , sham+analarmin (red open square symbols, n=14/4), mTBI+antalarmin (red filled square symbols, n=22/5); **p<0.01, ***p<0.001, ****p<0.0001 by Chi squared tests and 3-way ANOVA.
